# Temperature and depth dependence of the spatial distribution of snow crab

**DOI:** 10.1101/2022.12.20.520893

**Authors:** Jae S. Choi, Brent Cameron, Kate Christie, Amy Glass, Ellen MacEachern

## Abstract

The cascading effects of rapid climate change is a reality with which all biota are challenged. In this context, we examine the spatiotemporal probability of occurrence of snow crab as a means to express viable habitat. This is attempted for three demographic components, morphometrically mature males and females and immature adolescent crab in the Scotian Shelf region of the northwest Atlantic, Canada. We use a robust approach, known as Conditional AutoRegressive models, to define viable habitat. Further, we focus upon viable habitat, conditioned on the marginal influence of temperature and depth as they are known to be important constraints on snow crab. We observe some niche partitioning in terms of depth and temperature. We also note declines in viable habitat marginal to depth and temperature since 2010 for all demographic groups. This population representing the southern-most distribution of snow crab in the northwest Atlantic are vulnerable to degradation of viable habitat attributable to rapid climate change.

**One-Sentence Summary:** Rapid climate change and a decadal scale change in the viable habitat of snow crab of the Scotian Shelf ecosystem.

## Introduction

Temperature is a profoundly important component of the habitat of all organisms (Kleiber, 1961). Snow crab (*Chionoecetes opilio*, O. Fabricius) has a particularly narrow temperature preferendum (Elner and Bailey, 1986; Choi, 2011). Temperatures greater than 7°C are known to be detrimental to snow crab metabolic balance based upon laboratory studies (Foyle, O’Dor and Elner, 1989). There exist many other factors that can influence the viability of habitat (Elner and Bailey, 1986; Hooper, 1986; Comeau et al., 1998; Choi, 2011). Some, such as sediment type/granularity, depth, are more static features operating on very long time scales. Others include predator-prey and competitive interactions (Robichaud, Elner and Bailey, 1991; Boudreau, Anderson and Worm, 2011) as well as ocean acidification due to carbon geochemistry (Taylor et al., 2015). Depth in particular is a proxy for pressure, light levels, turbulence, substrate complexity, and overall variability of many environmental factors. In general, their interactions are not always straightforward and so there is usually a depth range that is preferred over others, depending upon the specifics of a region.

As temperature is also interlinked with many of these habitat factors, a model based approach is essential to elucidate its effect upon snow crab. For this purpose, we define “*viable habitat*” (see Methods and Materials) as a probability statement by using Bayesian spatiotemporal Conditionally Autoregressive (CAR) models that can borrow information from adjacent areas and times (Besag, York and Mollié, 1991; Rue, Martino and Chopin, 2009; Riebler et al., 2016; Brown and Zhou, 2021; Choi, 2022a). In this paper, viable habitat is further conditioned on the *marginal* influence of temperature and depth as they are known to be important constraints on snow crab.

Our focal area is the snow crab domain in the Scotian Shelf of Atlantic Canada (Figure 1) for which we have high quality survey information collected since the late 1990s to the present (Figure 2). The area is also the southernmost part of their spatial distribution in the northwest Atlantic and at the confluence of a number of different currents. As such, snow crab in the region are affected by highly variable ocean bottom temperatures (Drinkwater, 2005; Choi, 2011) and will be sensitive to large-scale warming trends in the region.

**Figure 1.**
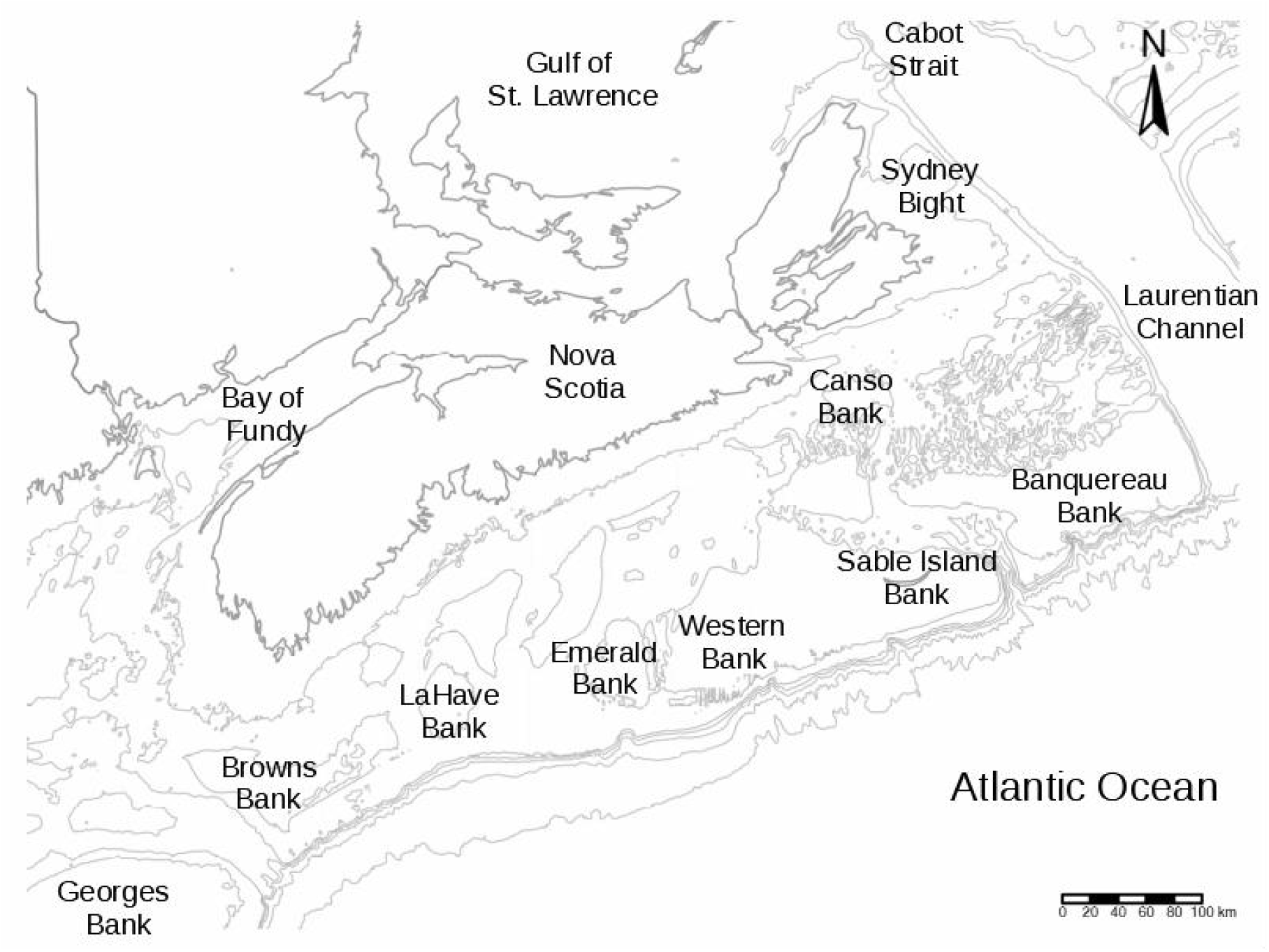
The area of interest in the Scotian Shelf of the northwest Atlantic Ocean (Canada, NAFO Div. 4VWX; UTM zone 20). Shown are isobaths (100, 200, …, 500, 1000 m) and some of the major bathymetric features in the area. This area is at the confluence of the Gulf Stream from the south and south east along the shelf edge, Labrador Current and St. Lawrence outflow from the north and north east, as well as a nearshore Nova Scotia current, running from the northeast. It is hydro-dynamically very complex due to mixing of cold, low salinity water from the north with the warm saline water from the south.

**Figure 2.**
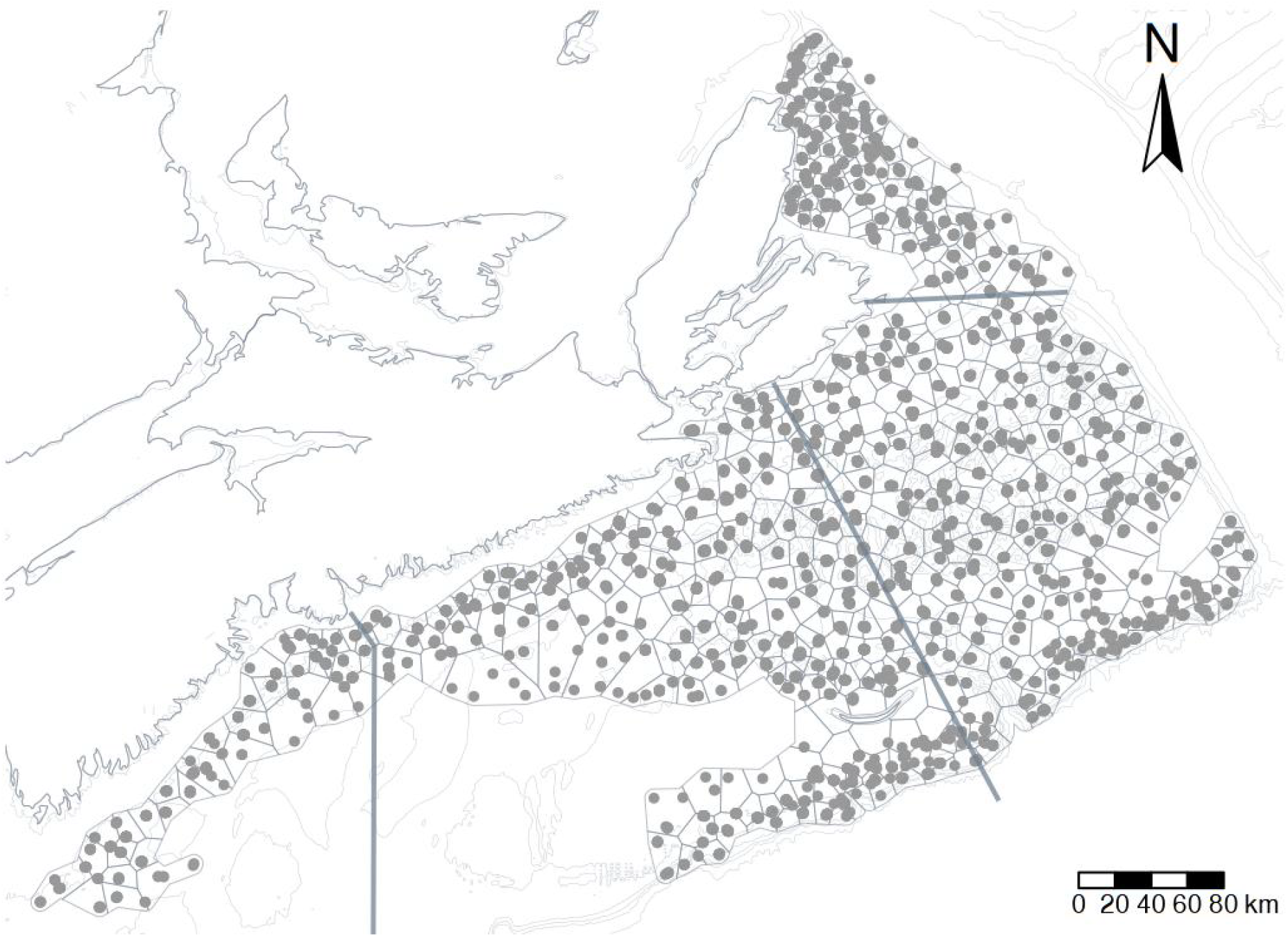
Spatial domain of snow crab survey locations (1999-2021). As most are fixed stations, each dot represents many sampling events. The spatial domain goes well beyond core fishing areas that are primarily deep cold-water basins. Snow crab are seldom found on sandy banks and so not sampled. Final estimation mesh is shown in fine solid lines. Thick solid lines delimit fishery management unit divisions,which are known as Crab Fishing Areas: 4X, 24, 23, and NENS (from left to right).

## Materials and Methods

Targeted annual surveys have been conducted in the region from 1999 to the present by Fisheries and Oceans Canada. A modified Bigouden Nephrops net designed to dig into sediments was used with sample tows of approximately 0.3 km in length and a target vessel speed of 3.7 km/h. Bottom contact and net configuration were monitored to provide explicit areal density estimates. The wing spreads ranged from 12 to 14 m, depending upon the substrate encountered. Supplemental information of multispecies associations were obtained from the Research Vessel surveys conducted annually by the Canadian Department of Fisheries and Oceans. The sampling stations for the latter were randomly allocated in depth-related strata with sample tows of approximately 1.82 to 2.73 km in length on a variety of different gears, mostly the Western IIA net. Vessel speeds ranged from 4.5 to 6.5 km/hr. Only tow-distances were available which was used to approximate areal density using a nominal wing spread of 12.5 m. In both surveys, all species are weighed and counted. In the RV surveys, subsamples are also individually measured.

We focus upon three demographic groups to verify the ontogenetic and sex-related partitioning of habitat noted in early observational studies (Robichaud, Bailey and Elner, 1989; Comeau and Conan, 1992; Sainte-Marie, 1993): immature crab, morphologically mature male crab, and morphologically mature female crab. Maturity is readily identifiable as snow crab have a terminal molt to maturity that results in a disproportionate increase in the size of the chela in males and abdominal flap in females, relative to overall carapace width. The onset of maturity is variable but usually occurs near 55 mm carapace width in females and 100 mm carapace width in males.

Specifically, we define “*viable habitat*” as a probability statement on the likelihood of observing a demographic group, in a given time and location, using the 0.05 empirical quantile of the non-zero numerical densities to discriminate between “*viable*” and “*non-viable*” habitat. As in (Choi, 2011) we categorize *viable habitat* as a *Bernoulli* process with a *logit* link function and conditional upon covariates (Royle and Dorazio, 2008). The spatial and temporal process Y_st_, are expressed at discrete areal units s=(1, …, a) and time units t=(1, …, b) with autocorrelated random effects in space, time and spacetime (Choi, 2022a):

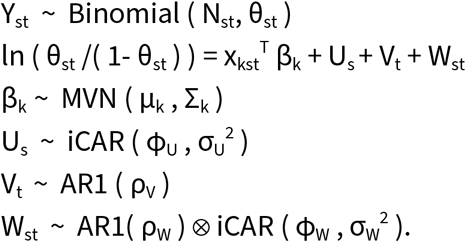

Here, θ_st_ is the vector of probabilities of success in each trial in each space and time unit and N_st_ is the number of trials; the covariates x_kst_^T^ reshaped to appropriate matrix form and associated covariate coefficients parameters β_k_ with a multivariate normal prior with mean μ_k_ and diagonal variance matrix Σ_k_; the spatial autocorrelation U_s_ follows the iCAR (also known as a convolution model or the Besag-York-Mollié or “BYM” model in the epidemiological literature, when data are Poisson distributed; (Besag, York and Mollié, 1991; Riebler et al., 2016; Lawson, 2018)) with ϕ_U_ the proportion of the iCAR variance, σ_U_^2^, attributable to spatial processes relative to nonspatial processes; the temporal error components V_t_ are assumed to follow a lag-1 autocorrelation process (AR1) with parameter ρ_V_; and finally, the spatiotemporal errors W_st_ are modeled as a Kronecker product (⊗) of temporal AR1 and spatial iCAR processes, where time is nested in space. This model form is known as an “inseparable” space-time model. The *k* covariates in this study were: bottom temperatures, depth, season, species composition ordination axes (1 and 2), and a global intercept term as the overall domain average for *viable habitat*. These are factors known apriori from the literature to be important to habitat selection of snow crab (see references in Introduction). Species composition was expressed using Principal Components Analysis of co-occurring species from snow crab surveys and groundfish research surveys after expressing their relative densities as z-scores. The first two axes represented approximately 13.2 and 7.7% of the total variance (145 species representing the 99.5% of the top occurrences in the region) and a simple means of parameterizing the community structure and potential ecological interactions.

Areal units were defined using an iterative Voronoi tessellation based algorithm that started with an ultrafine mesh, which was incrementally simplified by dropping areal units with low numbers of sampling stations (Choi, 2022b). This was continued until the station density distribution (no. per areal unit) was approximately Poisson (mean equal to variance), with a global maximum of 30 sampling events across all years within a given areal unit to permit identification of temporal patterns within a given areal unit; and total number of areal units of less than 2000 units (for computational stability and time). The approach permitted a spatial mesh with informative areal units capable of resolving temporal trends and environmental gradients. This was, therefore, a balance between information gain by maximally discriminating space and time and the costs of increased computational time and decreased model stability.

All analyses were conducted with R and R-INLA with algorithms and code available online (https://github.com/jae0/bio.snowcrab). Note that we utilize the “bym2” parametrization of the iCAR for stability and interpretability of parameter estimates (Riebler et al., 2016). Penalized Complexity (PC) priors that shrink towards a null (uninformative base model) were used for all random effects (Simpson et al., 2017). For the AR1 process (V_t_), we used the “cor0” PC prior which has a base model of 0 (i.e., no correlation). For the spatial process (“bym2”) we use the PC prior (rho=0.5, alpha=0.5; U_s_ and W_st_). Seasonality was discretized into 10 units and a cyclic “rw2” process was used. For all other covariates, a second order random walk process (“rw2”) was used as a smoother using 11 quantiles as discretization points.

## Results and Discussions

In the focal domain, an increase in annual mean predicted bottom temperatures has been evident from about 6°C prior to 2012, to almost 8°C in 2021 (Figure 3). Biologically, this is important, especially as it bounds the 7°C metabolic breakpoint. However, the spatiotemporal variability (that is, persistence of temperature effects) associated with this increase is also very large and so the biological relevance to snow crab needs to be clarified. The CAR model helps to disentangle the *realized* thermal preferences from other important factors. This clarification is accomplished by accounting for the spatial, temporal and spatiotemporal autocorrelations and features of interest, notably, depth and temperature as they are known in the literature to have a strong influence upon snow crab biology and ecology (Elner and Bailey, 1986; Robichaud, Bailey and Elner, 1989).

**Figure 3.**
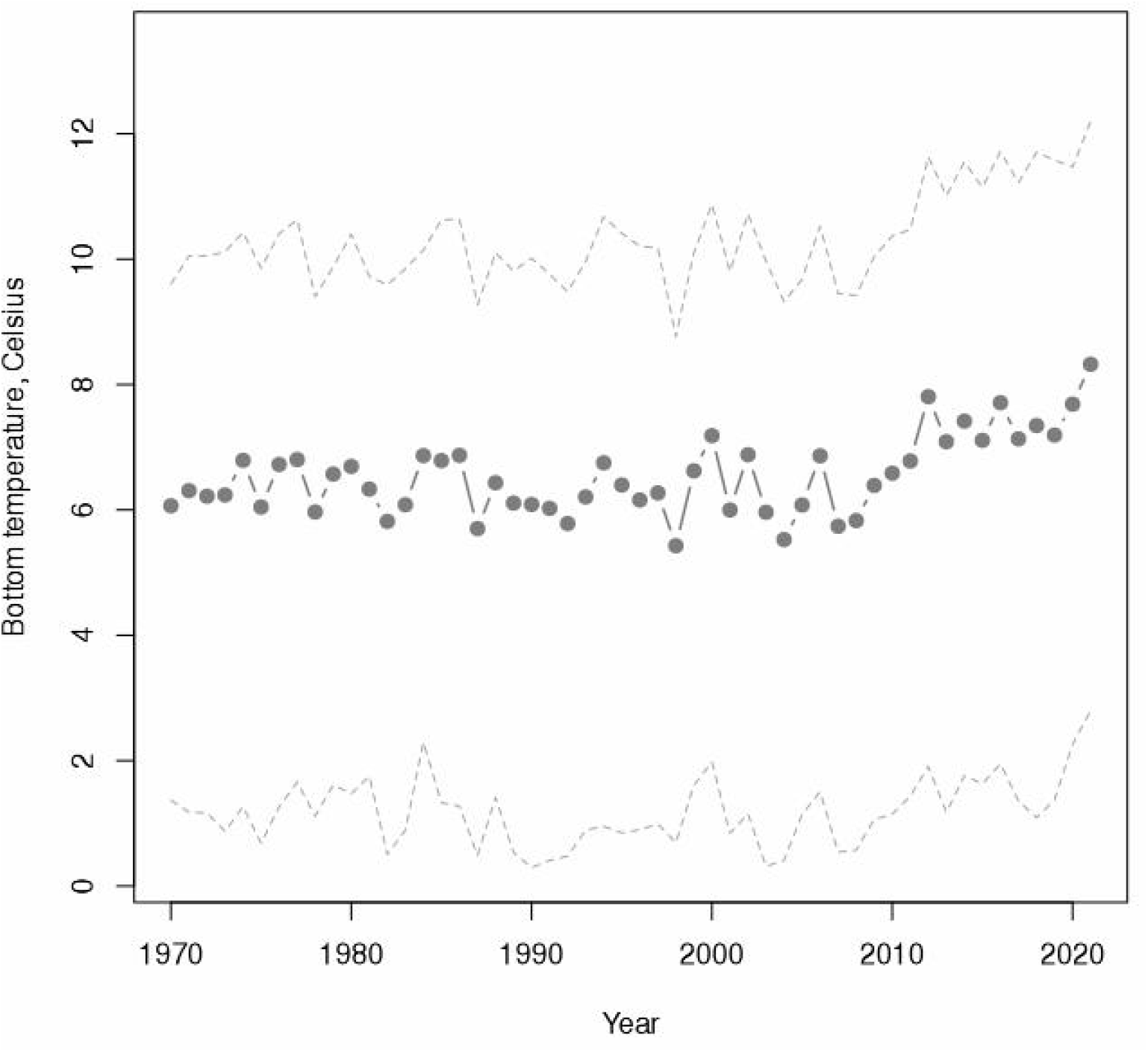
Posterior predicted mean bottom temperatures (°C) on the snow crab domain of the Scotian Shelf. Stippled lines are 95% Credible Intervals.

The overall posterior predictions of the temporal patterns of *viable habitat* of each demographic group was similar (Figure 4). A low was evident in 2004, though for immature crab this was a bit earlier, in 2002. Thereafter, *viable habitat* quality increased to a peak in 2007/2008 and then declined again to 2015/2016. After a short interval of increase they decreased in 2021 to historical lows. The exception being mature females that experienced an even lower overall *viable habitat* quality in 2004.

**Figure 4.**
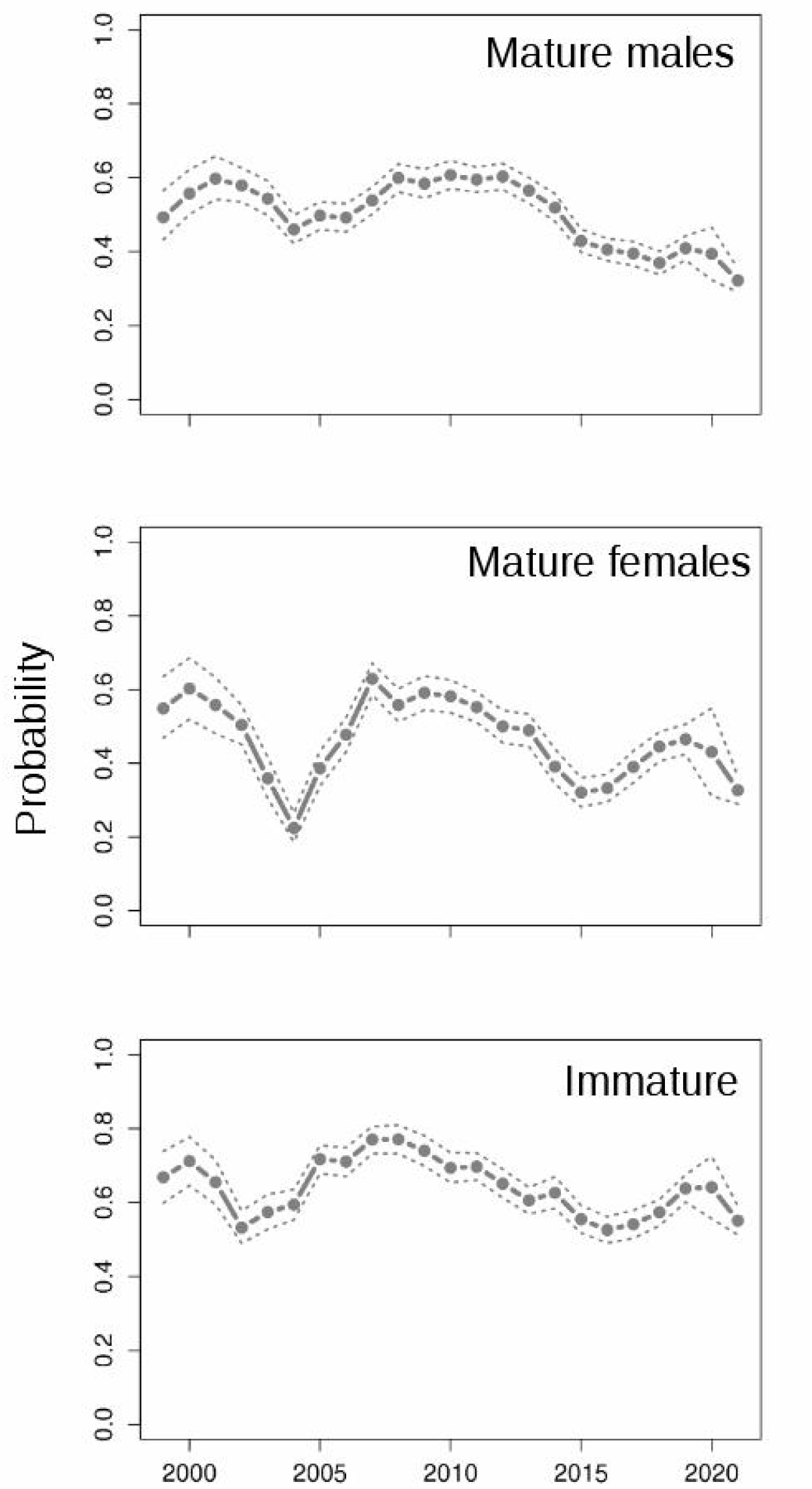
Posterior predicted overall mean *viable habitat* (probability) on the Scotian Shelf as a function of time. Stippled lines are 95% Credible Intervals.

The so-called “persistent” (mean) spatial effects provide an indication of core areas of *viable habitat* (Figure 5). Immature crab are found with higher probability closer to shore in bathymetrically more variable areas. Mature male snow crab are also found further offshore than immature snow crab in and along deep channels. Although there is overlap, a clear separation is evident in their spatial distributions. Female crab seem to have a more diffuse distribution that is somewhat intermediate to those of mature males and immature snow crab. No specific core area of recruitment or nursery areas were evident and instead the whole region seems to be utilized.

**Figure 5.**
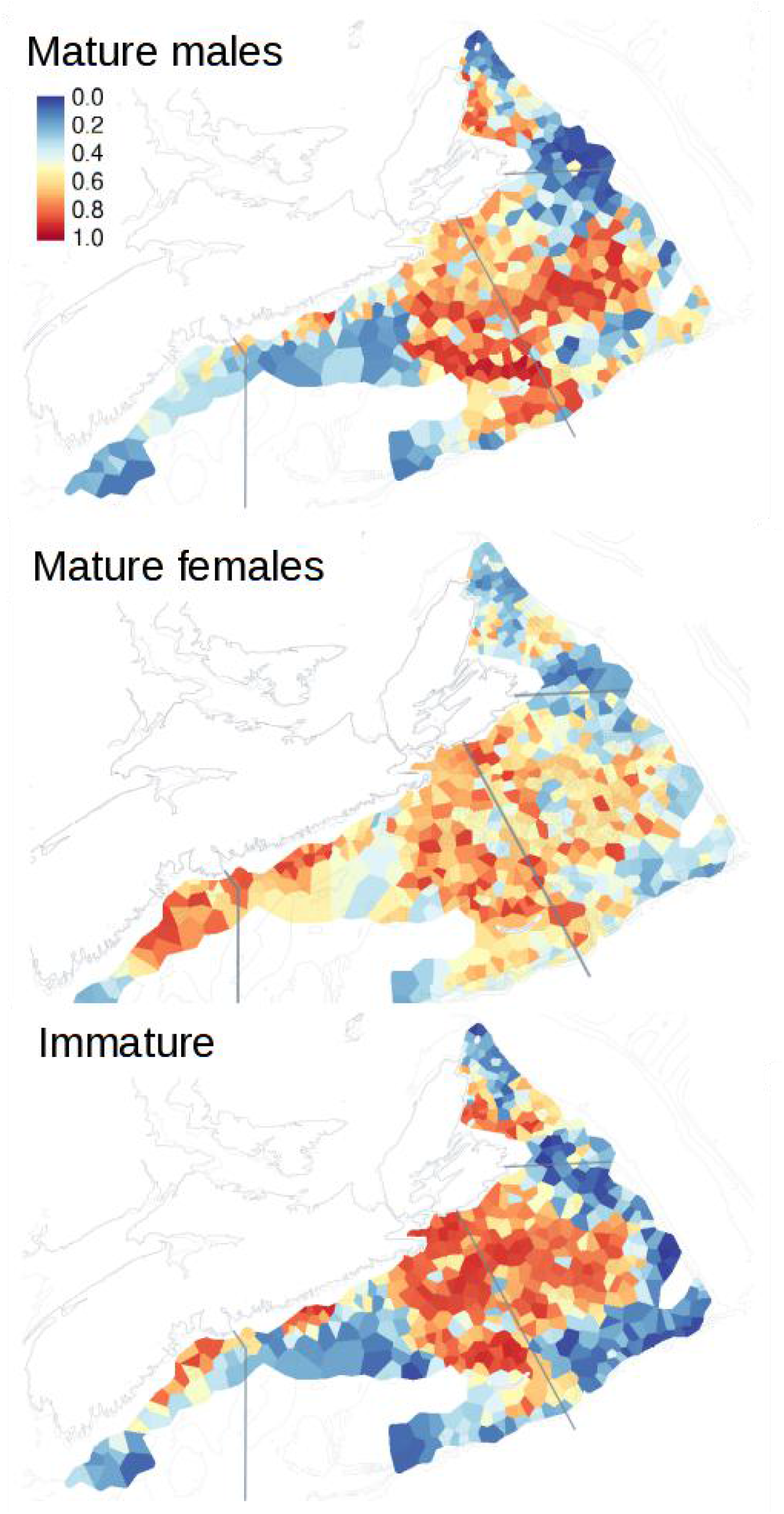
Posterior spatial effects on *viable habitat* (probability); that is, persistent spatial patterns or “core” habitat areas. The colour scale is on a probability scale.

The posterior predictions across time (Figure 4) are overall means and so convolved with variations in overall density associated with natural population cycles and persistent spatial effects. The *marginal* relationship of depth and temperature independent of these other factors are particularly informative in the context of rapid climate change (Figure 6). Using the 0.5 probability isocline as a heuristic landmark, depth and temperature related partitioning of their environment can be seen more clearly. Mature males had a preference of depths between 160 to 300 m and temperatures of 4°C or less. Mature females had a preference for shallower depths (100 to 160 m) and colder temperatures of 0°C or less. Immature snow crab preferred greater depths (> 160 m), similar to mature males, and temperatures of 3°C or less. Rapid climate change can reduce available viable habitat.

**Figure 6.**
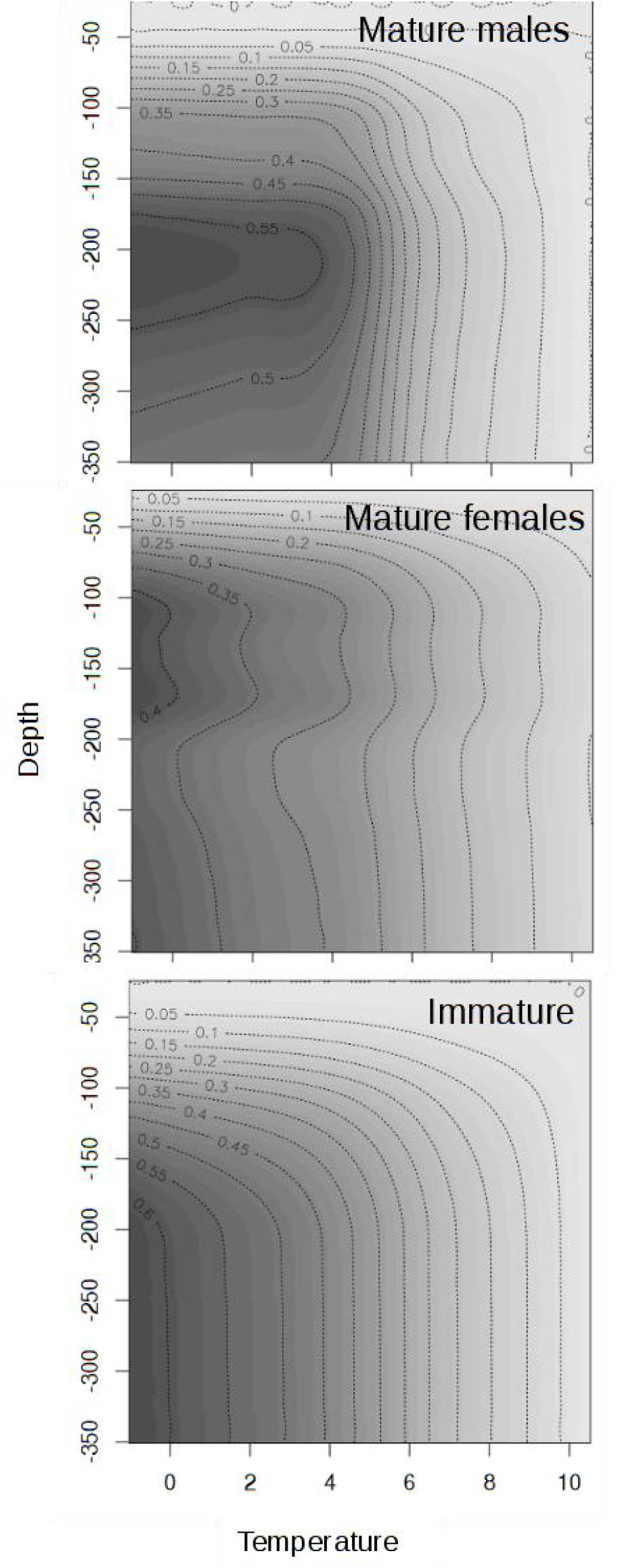
Posterior depth-temperature effects upon snow crab *viable habitat* (probability). Using the 0.5 probability isocline for comparisons: mature males prefer depths of 160 to 300 m and temperatures of 4 °C or less; mature females prefer 100 to 160 m and 0 °C or less; and immature snow crab prefer 160 to > 350 m and 3 °C or less.

Using these *marginal* effects of depth and temperature, we estimated the surface area associated with marginal probabilities greater than 0.25 for the above mentioned highest probability depths (Figure 7) as an indicator of changes in the amount of high-quality habitat area. The overall average surface area associated with these preferred depth ranges that intersect with preferred temperatures has varied significantly and coherently for each of the demographic groups. *Viable habitat* peaked in 1998 and then after a number of very large fluctuations, stabilized at a low level from 2012 to 2020. In 2021, they declined further to all time historical lows at levels of approximately 50% less than those of 1998, simply due to temperature-depth effects. The observed increase in temperature (Figure 3), though initially too variable to make any inference due to the large credible intervals, was nonetheless, likely to have been biologically significant once examined in the context of the recent declines of *viable habitat marginal* to depth and temperature (Figures 7) and the associated contraction of overall *viable habitat* for all three demographic groups (Figure 8). Other areas that experience similar environmental variability due to rapid climate variability need to be aware of and account for elevated mortality and spatial range contraction and distributional change.

**Figure 7.**
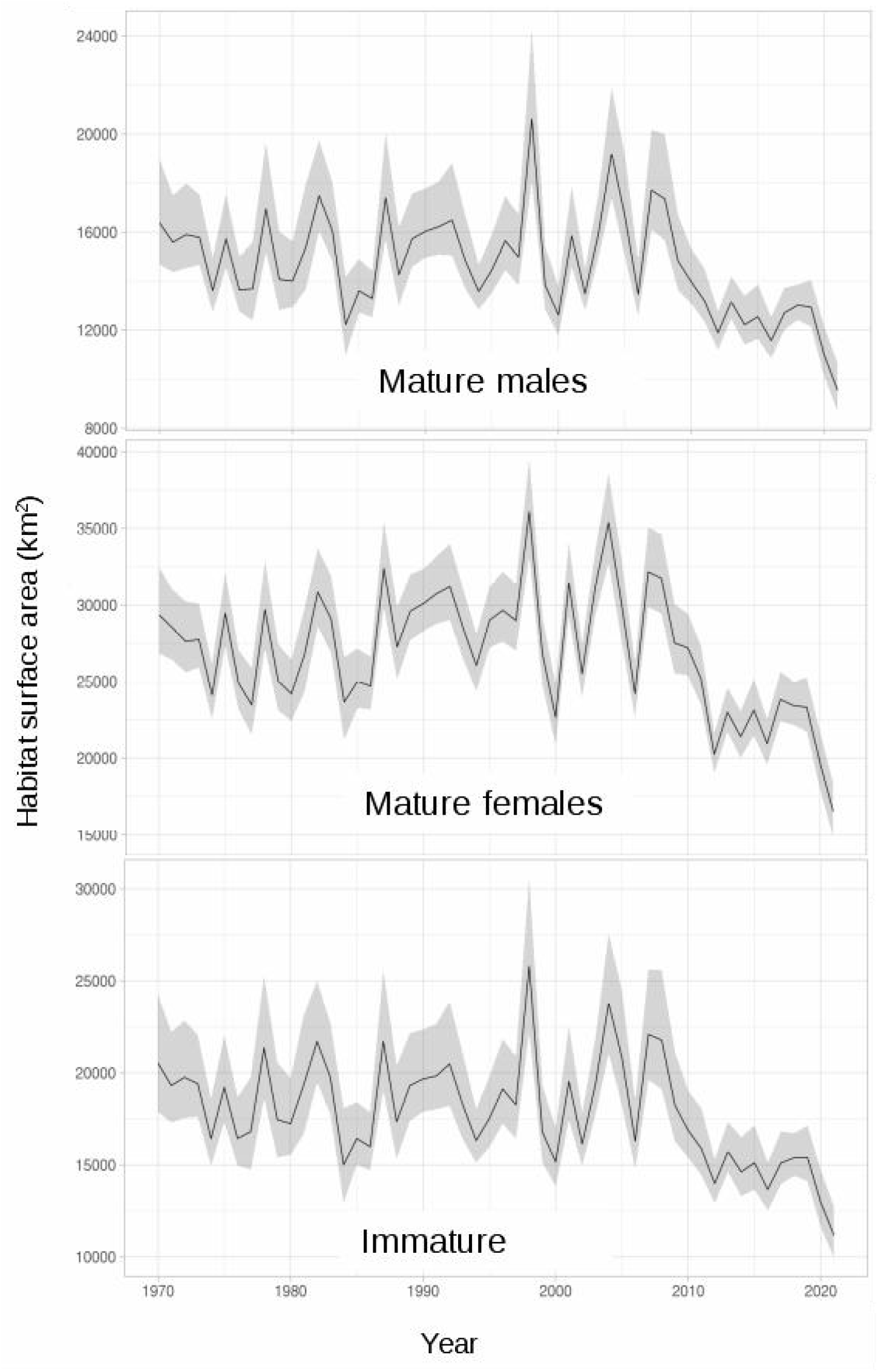
Posterior distribution of the surface area of *viable habitat marginal* to temperature and depth, with a marginal probability > 0.25. The preferred depth ranges are identified in the caption for Figure 6.

**Figure 8.**
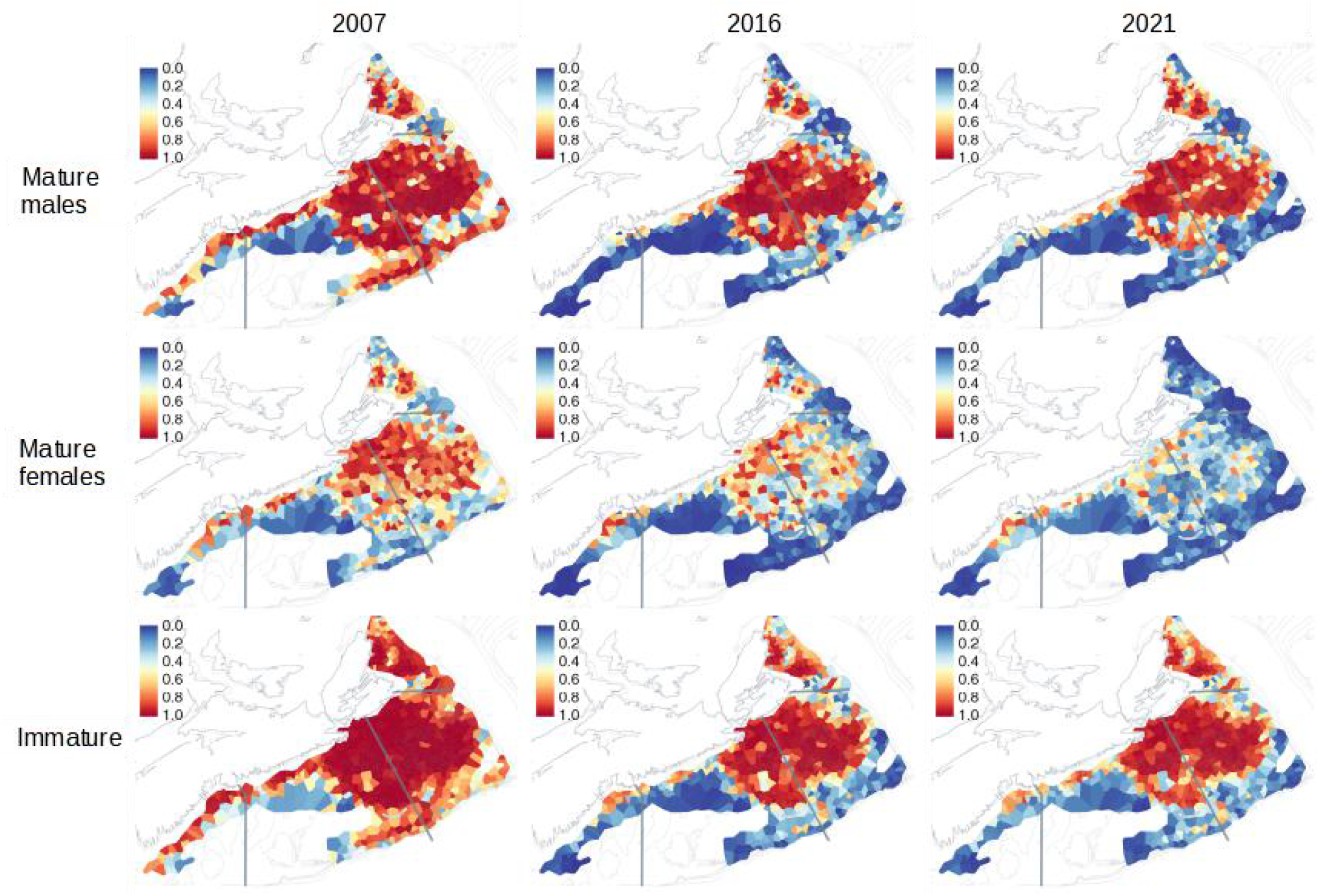
Map of posterior **predicted** *viable habitat* in key years. Year 2016 is an example of the overall low *viable habitat* conditions during the 2012-2020 period. Year 2007 is an example of a higher *viable habitat* just prior to this period and year 2021 is associated with the lowest *viable habitat* in the historical record.

## Conclusions

Snow crab, due to their narrow habitat preferences can be seen as a biological indicator of ocean integrating spatiotemporal variations in bottom temperature condition. Their distributions in the area of study suggests that the bottom climate has undergone some biologically significant and persistent changes that have lasted at least a decade. Even with such persistent changes, they continue to survive in the area. However, if poor conditions continue to persist or degrade further, there is a risk of extirpation from the area and a shift in spatial distributions to colder environments, further north.

## Scientific Reviews

Insightful and critical reviews were kindly provided by Drs. H. Bowlby and S. Boudreau.

## Data and materials availability

All data are held in databases in Fisheries and Oceans Canada. All analytical code are available at https://github.com/jae0/bio.snowcrab/.

We also acknowledge the efforts of numerous people that have collected and maintained this data over the years.

The perspectives outlined here are those of the authors and not necessarily those of funders nor Fisheries and Oceans Canada.

